# What NLR you recognizing? Predicted binding affinities- and energies may be used to identify novel NLR-effector interactions

**DOI:** 10.1101/2024.07.26.605369

**Authors:** Alicia Fick, Jacobus Lukas Marthinus Fick, Velushka Swart, Noëlani van den Berg

**Author notes:** Correspondence: Noëlani van den Berg University of Pretoria Department of Biochemistry, Genetics and Microbiology Pretoria, 0002, South Africa.

## Abstract

Nucleotide binding Leucine-rich repeat (NLR) proteins play a crucial role in effector recognition and activation of Effector triggered immunity in plants following pathogen infection. Advances in genome sequencing have led to the identification of a myriad *NLRs* in numerous agriculturally important plant species. However, deciphering which NLR proteins recognize specific pathogen effectors remains a challenge. Predicting NLR-effector interactions *in silico* would provide a more targeted approach for experimental validation, critical for elucidating function, and advancing our understanding of NLR-triggered immunity. In this study, NLR-effector protein complex structures were predicted using AlphaFold2-Multimer for all experimentally validated NLR-effector interactions. Binding affinities– and energies were predicted using 97 machine learning models from Area-affinity. We show that predicted structures with an AlphaFold confidence score > 0.42 have acceptable accuracy, and can be used to investigate NLR-effector interactions *in silico*. Binding affinities for 58 NLR-effector complexes ranged between –8.5 and –10.6 log(K), and binding energies between –11.8 and –14.4 kcal/mol, depending on the Area-Affinity model used. For 2427 “forced” NLR-effector complexes, these estimates showed larger variability, enabling the identification of novel NLR-effector complexes with 99% accuracy using an Ensemble machine learning model. The narrow range of binding energies– and affinities for true interactions suggest a specific change in Gibbs free energy, and thus conformational change, is required for NLR activation. This is the first study to provide a method for predicting NLR-effector interactions, applicable to all plant-pathogen interactions. Finally, the NLR-Effector Interaction Classification (NEIC) resource can streamline research efforts by identifying NLRs important for providing resistance against plant pathogens, advancing our understanding of plant immunity.

## Introduction

During pathogen infection of plants, both species produce numerous proteins that influence the outcome of infection (Ammari et al., 2016; Li et al., 2022; Naidoo et al., 2018). Plant nucleotide binding-leucine rich repeat (NLR) proteins recognize pathogen effectors secreted into the cytoplasm of host cells, subsequently activating host immune responses (Dodds et al., 2024; Tang et al., 2017). Effector proteins can interfere with the host immune signaling cascades, when undetected by NLRs, which results in successful pathogen infection (Zhang et al., 2022). Due to their critical role, NLRs are the focus of many plant-pathogen interaction studies (Contreras et al., 2023; de Araújo et al., 2019).

The increase in plant genome and RNA-sequence data has contributed to the identification of numerous *NLRs*, especially in important agricultural crops (Li et al., 2023; Liu et al., 2024; Santos et al., 2022; Yu et al., 2023). Furthermore, studies have successfully targeted NLR proteins for modification to increase resistance to pathogens (Contreras et al., 2023a; Maidment et al., 2023; Tamborski et al., 2023). However, the identification of the specific NLRs that recognize particular pathogen effectors, and activate immune responses thereafter, remains challenging due to the complexity of plant-pathogen interactions, and the multitude of possible NLR-effector interactions (Barragan et al., 2021; Yang et al., 2019; Zheng et al., 2021). Experimental validation like yeast two-hybrid systems or co-immunoprecipitation assays, are complex, experimentally cumbersome, and expensive (Zheng et al., 2021).

Studying NLR-effector interactions, and identifying functionally important NLRs, can be simplified by focusing on singleton NLRs, which directly recognize effectors, and activate immune responses (Adachi and Kamoun, 2022; Contreras et al., 2023). Other NLRs may form part of NLR networks, with sensor NLRs binding to effectors and activating helper NLRs to trigger immune response activation (Contreras et al., 2023), or they may guard effector-targeted proteins, without direct binding (Jones and Dangl, 2006). Identifying NLRs that directly bind to effectors would simplify predictions of unknown NLR-effector interactions. Of 419 experimentally activated NLRs, a total of 67 NLRs recognize 93 effectors directly (Brambam et al., 2024; Kourelis et al., 2021; Redkar et al., 2022).

Most data on NLRs are from the Asterids, which have highly diversified sensor– and helper NLR families (Adachi and Kamoun, 2022; Wu et al., 2017). Studies on NLRs in other plant groups such as Rosids, including NLRs from *Arabidopsis thaliana*, *Vitis vinifera* (grape), and *Manihot esculenta* (cassava), show a higher percentage of singleton NLRs with no network-related NLRs present in these genomes (Wu et al., 2017). Thus, it is expected that more singleton NLRs exist in the majority of species for which genomic sequences are not yet available. Prigozhin and Krasileva (2021) showed that direct recognition NLRs have higher amino acid diversity, per amino acid site – calculated with the use of Shannon entropy scores. Higher Shannon entropy scores were particularly observed within the Leucine-rich repeat (LRR) domains, which bind to effectors and govern recognition specificity. Thus, we hypothesize that Shannon entropy scores can predict singleton NLRs with direct effector recognition capability. Following the identification of direct recognition NLRs, predictions can be made regarding the specific effectors recognized by individual NLRs.

Various programs for predicting plant-pathogen protein-protein interactions are available (Lei et al., 2024; Li and Zhang, 2016; Yang et al., 2019), using different methods such as protein domains (Deng et al., 2002), sequence similarity (Matthews et al., 2001), gene co-expression, and protein structure (Zhang et al., 2012). Machine learning (ML) programs have advanced plant-pathogen protein-protein interaction predictions. However, models are primarily trained on data from *Arabidopsis* and *Oryza sativa* (rice) (Karan et al., 2023; Lei et al., 2024; Yang et al., 2019; Zheng et al., 2021). Due to NLR effector-recognition specificity being influenced by single-nucleotide mutations in both NLRs and effectors, current programs may be inadequate for predicting NLR-effector interactions in plant species without protein-protein interaction data (Dodds et al., 2001; Ortiz et al., 2022; Segretin et al., 2014; Tamborski et al., 2023). The limited availability of true NLR-effector interaction data further complicates predictions (Sharma, 2022).

In this study, we aimed to develop a method for predicting NLR-effector interactions *in silico*, to study plant-pathogen interactions. These predictions allow for a targeted approach to investigate both NLR– and pathogen effector functionality. Once more data are available for NLR-effector pairs, dedicated ML programs can be developed to analyze plant-pathogen protein-protein interactions more accurately. We demonstrate that NLR-effector protein structures can be predicted by AlphaFold2-Multimer with acceptable accuracy, comparable to experimentally validated structures, enabling their use in studying NLR-effector interactions. Predicted structures were used to determine binding affinities (BA) and binding energies (BE) using multiple ML models, for which data were combined and used for the training of an Ensemble learning model. Differences in BA– and BE values were observed between true NLR-effector partners, and forced NLR-effector partners (known not to activate plant immune responses), allowing the prediction of interaction probability using the trained Ensemble model. These results enhance our understanding of NLR-effector interactions and NLR activation.

## Materials and Methods

### NLR and effector sequences

NLR and effector protein sequences, from 93 known interactions previously identified by Kourelis et al. (2021), were downloaded from the Uniprot online database (https://www.uniprot.org). NLR sequences included those from barley (*Hordeum vulgare* subsp. *vulgare*), bread wheat (*Triticum aestivum*), crab apple (*Malus* x *robusta*), tomato (*Solanum lycopersicum*), muskmelon (*Cucumis melo*), Arabidopsis (*A. thaliana*), Indica rice (*O. sativa* sub sp*. indica*), Japonica rice (*O. sativa* sub sp*. japonica*), flax (*Linum usitatissimum*), potato (*Solanum tuberosum*, *Solanum bulbocastanum*, *Solanum x edinense*, *Solanum venturi*, *Solanum chacoense*), soybean (*Glycine max*), tobacco (*Nicotiana benthamiana*), cultivated einkorn wheat (*Triticum monococcum* sub sp. *monococcum*), rye (*Secale cereale*), wild chilli pepper (*Capsicum chacoense*), and maize (*Zea mays*). Additionally, NLR sequences from *S. chacoense* (Rpi-chc1.2_543-5; QZA82918.1) and *S. tuberosum* (rpi-tub1.3_RH89-039-16; QZA82920.1) were downloaded from Uniprot. Amino acids in these sequences were manually altered according to experimentally tested mutations from Monino-Lopez et al., (2021). Three mutated versions of these NLR^LRR^ structures were used.

Pathogen effector sequences included oomycete effectors (*Hyaloperonospora arabidopsidis*; *Phytophthora infestans*; *Phytophthora sojae*), fungal effectors (*Blumeria graminis* f. sp. *hordei*; *B. graminis* f. sp. *secalis*; *B. graminis* f. sp. *triticale*; *B. graminis* f. sp. *tritici*; *Fusarium oxysporum* f. sp. *lycopersici*; *F. oxysporum* f. sp. *melonis*; *Magnaporthe oryzae*; *Melampsora lini*; *Parastagonospora nodorum*; *Pyrenophora tritici-repentis*) and bacterial effectors (*Erwinia amylovora*; *Pseudomonas syringae*; *Ralstonia pseudosolanacearum*; *Xanthomonas campestris* pv. *vesicatoria*; *Xanthomonas oryzae* pv. *oryzae*; *Burkholderia andropogonis*). Additionally, two *P. infestans* PexRD31 sequences (XP_002897637 and XP_002897638) were used for investigating interactions with Rpi-chc1.2 and rpi-tub1.3 (Monino-Lopez et al., 2021).

Leucine-rich repeat domains of NLRs (NLR^LRR^)– and effector signal domains were identified using the online InterPro web tool with default parameters. For protein prediction, only the NLR^LRR^ domain sequences were used, as these domains are the main determinants for effector recognition specificity in singleton NLRs (Kourelis et al., 2021; Locci and Parker, 2024). Effector signal domains were removed before further analysis. The complete dataset used is available in the Supplementary Data.

### NLR^LRR^-effector protein complex prediction

Protein complex structures were predicted using AlphaFold2-Multimer v.3 using Google Colab (Accessed September 2023 – July 2024), utilizing PDB100 templates and unpaired multi-sequence alignments (MSA) (Evans et al., 2021). Complex structures for all known interacting NLR^LRR^-effector partners (93 complexes) were predicted, as well as forced NLR^LRR^-effector complexes (5930 complexes). For each complex, five structures were predicted, and the structure with the highest pLLDT (per-residue model confidence score), pTM (predicted template modeling score), ipTM (interface pTM), and AlphaFold (AF) confidence scores (0.8*ipTM+0.2*pTM) was selected for further analysis (Evans et al., 2021). An AF confidence score > 0.5 indicates acceptable model accuracy; however, accurate protein structures with lower AF confidence scores have also been observed (Bret et al., 2024; Johansson-Åkhe and Wallner, 2022; Si and Yan, 2024; Yin and Pierce, 2023). Therefore, it is necessary to determine a relevant AF confidence score cut-off value to identify accurate NLR-effector complexes.

### Assessment of NLR^LRR^-effector prediction accuracy

The accuracy of the predicted structures was assessed by comparing them with available cryogenic-electron microscopy (cryo-EM) structures. NLR dimer and NLR integrated domain cryo-EM structures were excluded from these comparisons. Cryo-EM structures used for analysis included Sr35-AvrSr35 (7XVG), RPP1-ATR1 (7CRB), and Roq1-XopQ (7JLU), all downloaded from RCSB Protein Data Bank (https://www.rcsb.org; Ma et al., 2020; Martin et al., 2020; Zhao et al., 2022). Structures for these three NLR-effector complexes were predicted both with and without the use of PDB100 templates, using paired, unpaired, and unpaired-paired MSA options to intentionally introduce errors during the prediction process. This allowed for the comparison of structures with variable AF confidence scores to that of demonstrated cryo-EM structures. Five structures were predicted for each scenario, resulting in a total of 90 structures.

The predicted structures were compared with their respective cryo-EM structures using Dockground CAPRI-Q to obtain DockQ– and TM-scores, Critical Assessment of Predicted Interactions (CAPRI) rankings, and root mean squared difference (RMSD) values for both the backbone and sidechain interface residues (Collins et al., 2022; Basu and Wallner, 2016; Zhang and Skolnick, 2005). DockQ scores > 0.23 and TM-scores > 0.6 were used to indicate acceptable model confidence. Additionally, models were ranked based on the CAPRI criteria for protein-protein complexes (Lensik et al., 2017, Lensik and Wodak, 2010). These criteria include: High accuracy – Fraction of native contacts (fnat) ≥ 0.5, Ligand Root mean square (L-RMS) ≤ 1 and Interface Root mean square (I-RMS) ≤ 1; Medium accuracy – fnat ≥ 0.5, L-RMS > 1 and I-RMS > 1, or 0.3 ≤ fnat ≤ 0.5, L-RMS ≤ 5 or I-RMS ≤ 2; Acceptable accuracy – fnat ≥ 0.3, L-RMS > 5 and I-RMS > 2, or 0.1 ≤ fnat ≤ 0.3, and L-RMS ≤ 10, or I-RMS ≤ 4; Incorrect – fnat < 1, or L-RMS > 10 and I-RMS > 4. Following DockQ analysis and CAPRI ranking, a cut-off confidence score was determined to identify NLR-effector structures with acceptable accuracy.

The predicted complexes were visualized using ChimeraX v. 1.5. Interacting amino acids (AA) between the NLR^LRR^ and effector were identified using the “contacts” command, with any AA within 8 Å considered an interaction (UCSF ChimeraX; Meng et al., 2023; Pettersen et al., 2021). Hydrogen bonds were identified using the “hbonds” command, with a distance tolerance of 2.5 Å. The specific residue numbers involved in hydrogen bond formation were obtained and used for comparison between the predicted– and cryo-EM structures. In cases where residue numbers did not match between the two structures, the distance from the original interacting residue was calculated in both the N– and C terminal directions.

### Prediction of binding affinities– and energies for true and forced NLR-effector partners

The Area-Affinity online tool (accessed December 2023 – July 2024) was used to predict BA and BE (also known as Gibbs free energy) for all predicted NLR^LRR^-effector complex structures with an AF confidence score > 0.42. Predictions were made using both the protein-protein and antibody-protein programs from Area-Affinity (Yang et al., 2023). These programs collectively implement 97 different ML models for both BA and BE prediction, including linear, non-linear (random forest and neural network), constructed non-linear, generated non-linear, and mixed models. Results from all models were used to compare BA and BE values between true and forced NLR^LRR^-effector complex structures with AF confidence scores > 0.42. Area-Affinity was chosen for its superior or comparable accuracy relative to other predictive programs such as PRODIGY and Lisa (Raucci et al., 2018; Xue et al., 2016; Yang et al., 2022; Yang et al., 2023). Due to the limited availability of experimentally validated data for NLR-effector BEs and BAs, it was decided to use a combination of both protein-protein and antibody-protein models for BE– and BA prediction.

### Training of Ensemble learning model

An Ensemble classification model was developed to identify NLR^LRR^-effector interactions using all predicted BA– and BE values from the combined 194 Area-Affinity models. This Ensemble learning approach aims to combine informative data to improve predictive performance (Dong et al., 2020; Hastie et al., 2008). The model was implemented using the MATLAB Classification Toolbox v.24.1.0.2603908 (https://www.mathworks.com) and was trained with predicted BA and BE values for true and forced NLR^LRR^-effector complex structures with AF confidence scores > 0.42. The Classification Toolbox recommended Ensemble Boosted Trees with an Adaptive Boost model, which produced the most accurate results. This model combines several weak decision tree learners to form a strong classification algorithm (Mienye and Sun, 2022). The classifier model was trained to classify interactions as either true or forced, making the data labels binary with only two possible outcomes. Given the limited dataset available for classifier training, which could negatively affect the accuracy of more complex models, the use of Ensemble Boosted Trees with Adaptive Boost model was found to be an appropriate and effective solution.

It was found that BA and BE values from certain Area-Affinity models contributed more to the overall accuracy of the Ensemble model. Consequently, the ReliefF algorithm was used to select key features within the dataset (Kononenko et al., 1997). ReliefF uses a heuristic approach to determine the quality of the features and ranks them accordingly. Using the ReliefF algorithm, the top 21 features were identified as key contributors that diversify the dataset. Combining the Ensemble Boosted Trees using Adaptive Boost model with the ReliefF algorithm, greatly increased the model’s accuracy. The classification accuracy was determined using cross-validation with a KFold of 15. The trained model was then exported as an NLR^LRR^-effector interaction classification application (NEIC; available at https://github.com/koosfick/NEIC1.0). This application, created using MATLAB App Designer and MATLAB Compiler, utilizes the trained model and the ReliefF algorithm to classify NLR^LRR^-effector interactions from new input data.

### Statistical analysis and graphics

All statistical analyses were performed using RStudio v. 1.4.1106 (RStudio Team, 2020), and graphs were produced using the online Plotly web server (https://chart-studio.plotly.com).

The experimental procedure can be visualized in Figure 1.

**Figure 1.**
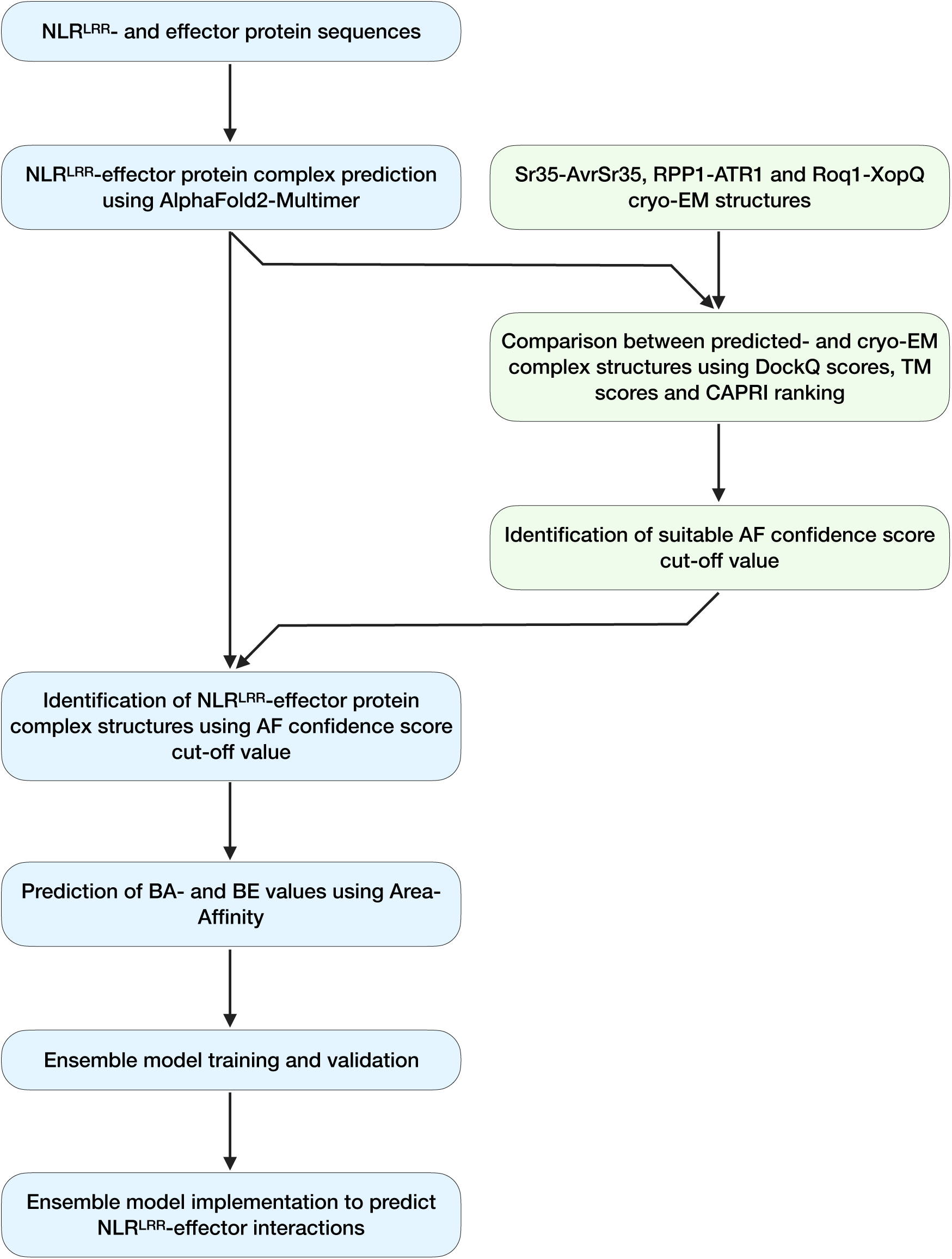
Visual representation of the experimental procedure followed in this study. Protein sequences were obtained from UniProt, with Leucine-rich repeat domains (LRRs) and effector sequences for protein structure prediction in AlphaFold2-Multimer. For effector sequences, signal domain-encoding sequences were removed. The accuracy of predictions and the determination of an acceptable AlphaFold (AF) confidence score cut-off value were assessed through comparisons with cryogenic electron microscopy (cryo-EM) structures. Once an AF confidence score cut-off value was established, Binding affinity (BA) and Binding energy (BE) values for all accurate NLR^LRR^-effector complexes were predicted using Area-Affinity. These values were used to train an Ensemble learning model, which was then implemented to classify NLR^LRR^-effector interactions as “true” or “forced”.

## Results and Discussion

### NLR-effector interaction prediction using a protein structure-based approach

Although ML programs have been developed to predict plant protein-protein interactions using a structure-based approach, these programs do not allow users to train new models on specific datasets (Dodds et al., 2001; Ortiz et al., 2022; Segretin et al., 2014; Tamborski et al., 2023). Moreover, the limited dataset for NLR-effector interactions may influence the accuracy of predictions made regarding these protein interactions. Therefore, we aimed to develop a novel method for predicting these interactions, by specifically focusing on comparisons between NLR^LRR^-effector data from true and forced interactions. First, we assessed the ability of AlphaFold2-Multimer to accurately predict structures of NLR-effector complexes and to identify an appropriate AF confidence score (Evans et al., 2021). Three cryo-EM structures of NLR^LRR^-effector complexes (Sr35-AvrSr35, RPP1-ATR1, and Roq1-XopQ) were compared with predicted complex structures having varying AF confidence scores (Ma et al., 2020; Martin et al., 2020; Zhao et al., 2022). A strong correlation between AF confidence scores and DockQ scores (R = 9.3, *p* < 0.01), as well as TM-scores (R = 0.85, *p* < 0.01) was observed, indicating that AF confidence can serve as an indication of complex prediction accuracy (Figure 2). Based on these metrics, an AF confidence score ≥ 0.42 was selected as an appropriate cut-off value for identifying predicted NLR^LRR^-effector structures with acceptable accuracy. Additionally, CAPRI-ranked models of medium quality also had AF confidence scores ≥ 0.4, supporting the validity of this cut-off value.

**Figure 2.**
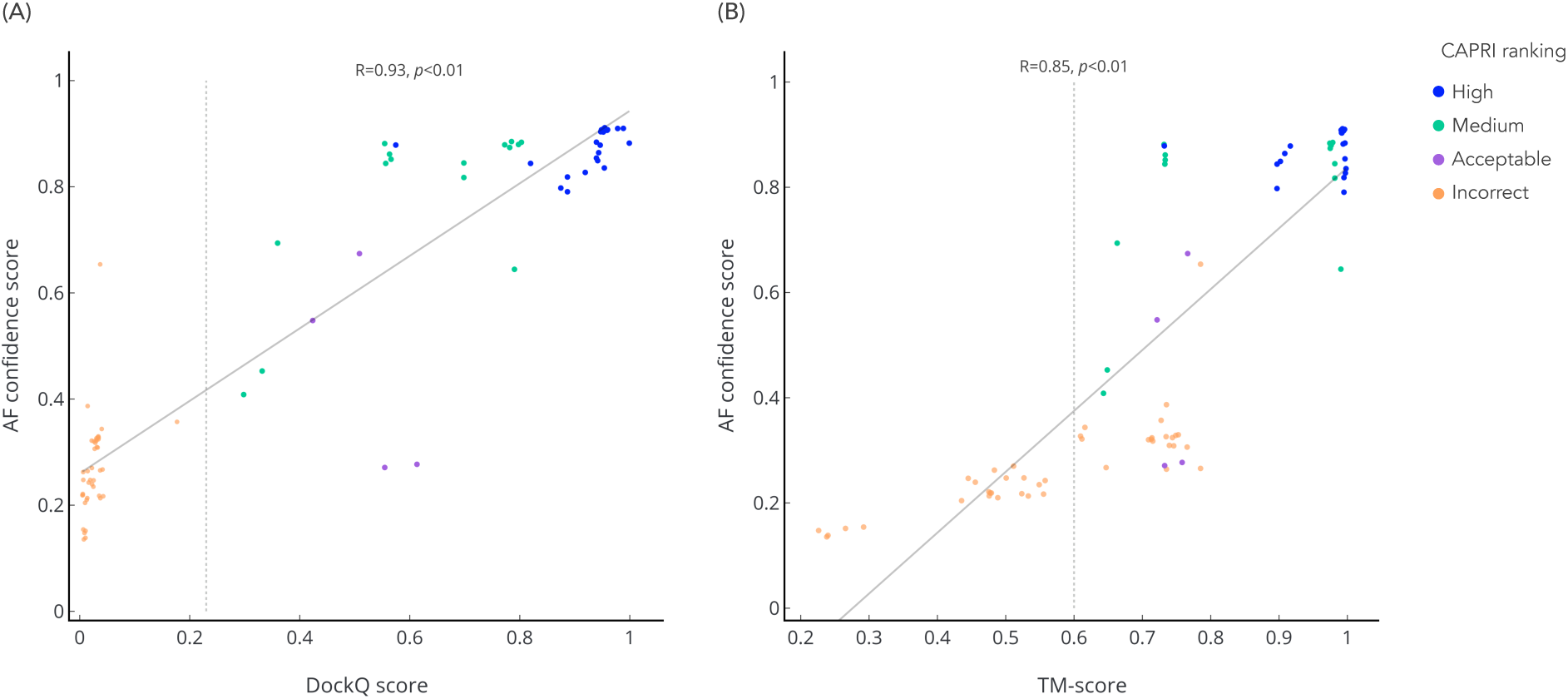
Model accuracy determined by the correlation between AlphaFold (AF) confidence scores, DockQ– and TM-scores. The distribution of AF confidence scores (calculated as 0.2*pTM+0.8*ipTM) for 90 predicted NLR^LRR^-effector structures is shown relative to **(A)** DockQ scores and **(B)** TM-scores. Structures for Sr35-AvrSr35, RPP1-ATR1, and Roq1-XopQ complexes were predicted using different protocols in AlphaFold2-Multimer v.3 and compared with native cryo-EM structures. Models were also ranked according to the stringent protein-protein complex criteria defined by the CAPRI community. Using DockQ– and TM-scores > 0.23 and > 0.6, respectively, an AF confidence score ≥ 0.42 was identified as indicative of acceptable model and model interface accuracy (PDB structure IDs: 7XVG, 7CRB, 7JLU). The Pearson’s correlation coefficient was used to measure the strength of correlation.

As expected, a negative correlation was observed between AF confidence scores and i-RMSDbb scores (R = –0.89, *p* < 0.01) and i-RMSDsc (R = –0.91, *p* < 0.1) (Figure 3). Using an RMSD value of < 3 Å for acceptability, 33 predicted structures with an average AF confidence score of 0.84 were identified (Manandhar et al., 2022; Ramírez and Caballero, 2018). Thus, a high AF confidence score is essential when investigating the specific interacting AA between NLRs^LRR^ and effectors *in silico*. Comparing the pattern of hydrogen bonds between predicted and cryo-EM NLR^LRR^-effector structures further highlights significant inaccuracies in the predicted interacting AAs (Figure 4A-C). These inaccuracies were particularly evident for effector proteins, with greater variation observed in the specific interacting AA when comparing predicted structures (with different AF confidence scores) to cryo-EM structures of the same NLR^LRR^-effector type. This is unsurprising, as effectors are often positioned within the concave structure of the LRR domain during interaction (Ma et al., 2020; Martin et al., 2020; Zhao et al., 2022). This restricts the interacting AAs of an NLR^LRR^ to the concave side of the LRR domain, while the effector can assume significantly more positions and orientations within the complex.

**Figure 3.**
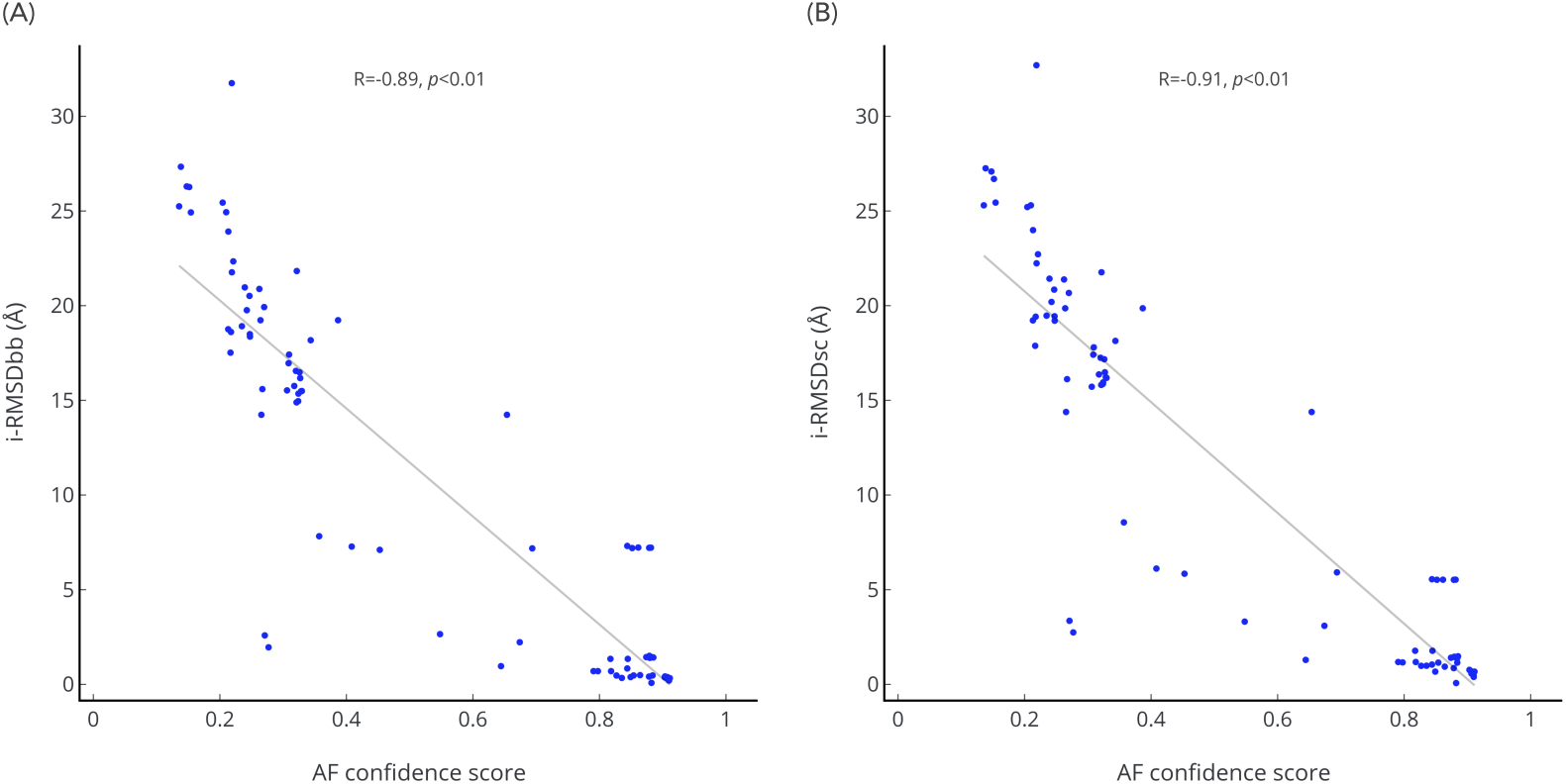
Correlation between AlphaFold (AF) confidence scores and interface Root Mean Squared Difference Scores (RMSD) scores for (A) interface backbone (i-RMSDbb) and (B) interface sidechain (i-RMSDsc) residues. The scores were obtained by comparing 90 predicted NLR^LRR^-effector structures (30 Sr35-AvrSr35 complexes, 30 RPP1-ATR1 complexes, and 30 Roq1-XopQ complexes) with cryogenic-electron microscopy structures. Structures were predicted using AlphaFold2-Multimer, following different protocols. AF confidence scores were calculated using pTM– and ipTM values (0.2*pTM+0.8ipTM). Pearson’s correlation coefficient values are shown above each graph. (PDB structure IDs: 7XVG, 7CRB, 7JLU).

**Figure 4.**
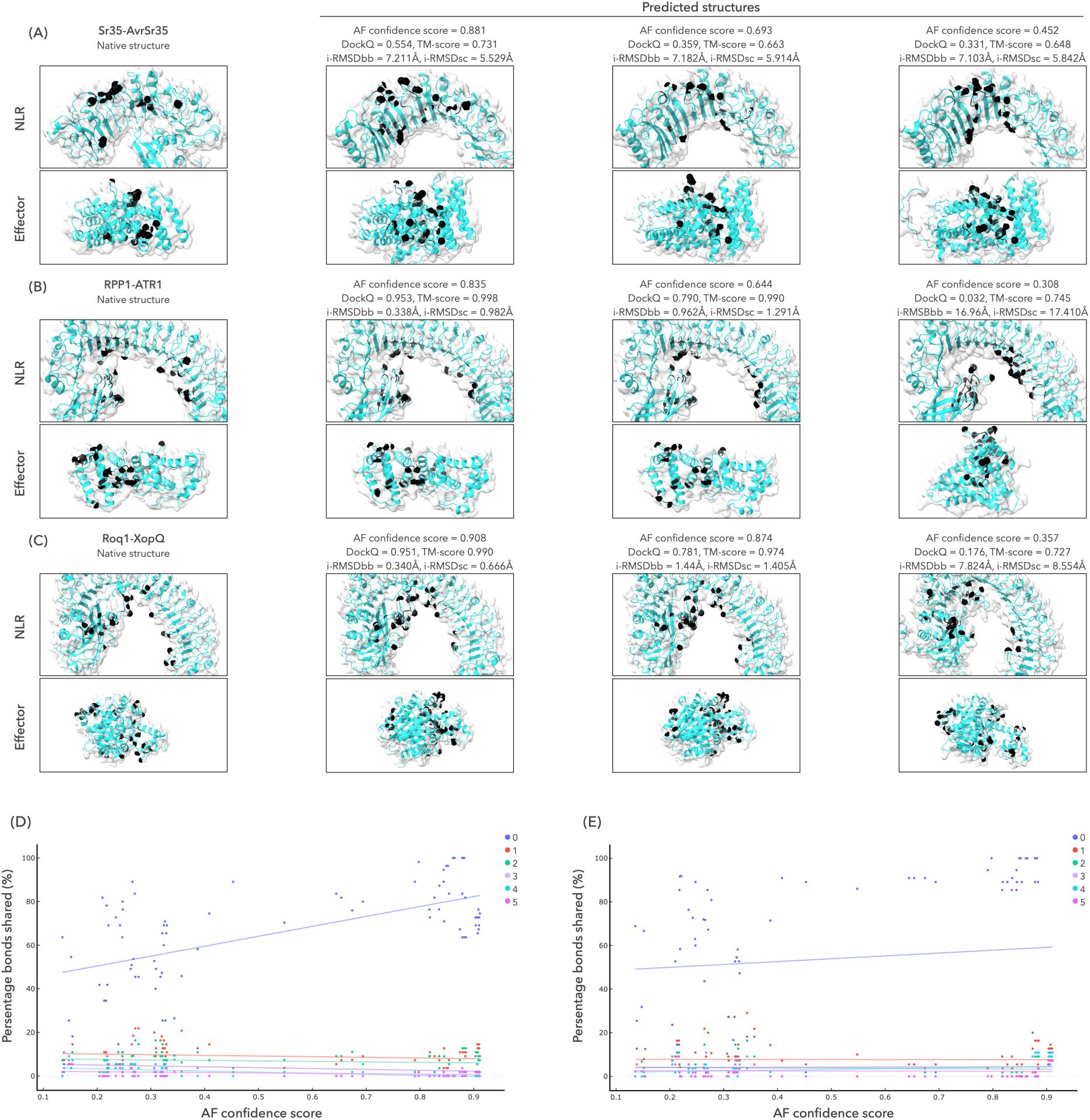
Location of amino acids (AAs) which participate in the formation of hydrogen bonds during NLR^LRR^ and effector interaction. Cryogenic-electron microscopy structures (native) and three representative AlphaFold2-Multimer predicted structures for **(A)** Sr35-AvrSr35, **(B)** RPP1-ATR1, and **(C)** Roq1-XopQ are shown. Each protein structure from an NLR^LRR^-effector pair is displayed separately, with NLR^LRR^ structures above effector structures. The AAs forming hydrogen bonds during NLR^LRR^-effector interactions are depicted in black. For predicted complexes, AlphaFold (AF) confidence scores (0.2*pTM+0.8*ipTM) are provided, along with DockQ– and TM-scores, and RMSD scores for interface backbone (i-RMSDbb) and interface sidechain (i-RMSDsc) residues. These scores were obtained through comparisons with the respective native structure. The percentage of shared AAs when compared between native and predicted structures of NLR^LRR^ **(D)** and effector **(E)** complexes with varying AF confidence scores is shown. The percentage is relative to the total amount of hydrogen bonds within the respective predicted NLR^LRR^-effector structure. AAs involved in hydrogen bond formation that are not shared with specific AAs in native structures are indicated when these AAs are 1 (red), 2 (turquoise), 3 (purple), 4 (blue), or 5 (pink) AA away (either in the N-or C-terminus direction) from the original bond location in the native structure. A significant correlation (R = 0.62; *p* < 0.01) was observed between the number of shared bonds for NLR^LRR^ structures, but no significant correlation was observed for bonds formed with effector proteins (R = 0.11; *p* > 0.05)

However, it is worth noting that an average of 80% interacting AAs are shared between predicted NLR^LRR^ structures with AF confidence scores > 0.6 and their corresponding cryo-EM structures. Additionally, less than 20% of predicted interacting AA are within 1-5 AA away from an interacting AA in the related cryo-EM structure (Figure 4D). As seen in Figure 4A-C, hydrogen bonds (depicted in black) are present within similar regions of predicted NLR^LRR^ domains, compared to the cryo-EM structures. These results indicate that a predicted NLR^LRR^-effector complex with an AF confidence score > 0.6 can be used to identify AAs that form bonds with the recognized effector protein. For predicted structures with AF confidence scores < 0.6, the average of shared interacting AAs decreases significantly to 53% (R = 0.62; *p* < 0.01), with more interacting AAs being predicted to be located further from interacting AAs in cryo-EM structures. Thus, predicted models with lower AF confidence scores may still be informative for *in silico* investigations of interacting AAs, albeit with limitations. This trend was not observed for interacting AAs relative to the effector protein, with no significant difference in the percentage of shared bonds between predicted and cryo-EM structures (R = 0.11; *p*> 0.05) (Figure 4E). Therefore, bonds between AAs in relation to the effector protein cannot be reliably used to predict which AAs have a higher probability of contributing to effector recognition by an NLR^LRR^.

### Comparisons between true– and forced NLR^LRR^-effector interactions

Since NLR^LRR^-effector structures with AF confidence scores > 0.42 were shown to have acceptable accuracy, we compared structures of true and forced NLR^LRR^-effector interactions to assess any identifiable differences between these two classes. In total, 6022 structures were predicted, including 93 true NLR^LRR^-effector interactions. Neither model nor interface accuracy (plDDT, pTM, and ipTM values) differed significantly between true and forced interaction complexes (Figure 5). Thus, contrary to a previous study by Martin (2024), high ipTM values cannot be used to distinguish between true and forced interactions. Instead, they only indicate a high confidence level of the interface position within the NLR^LRR^ structure (O’Reilly et al., 2023).

**Figure 5.**
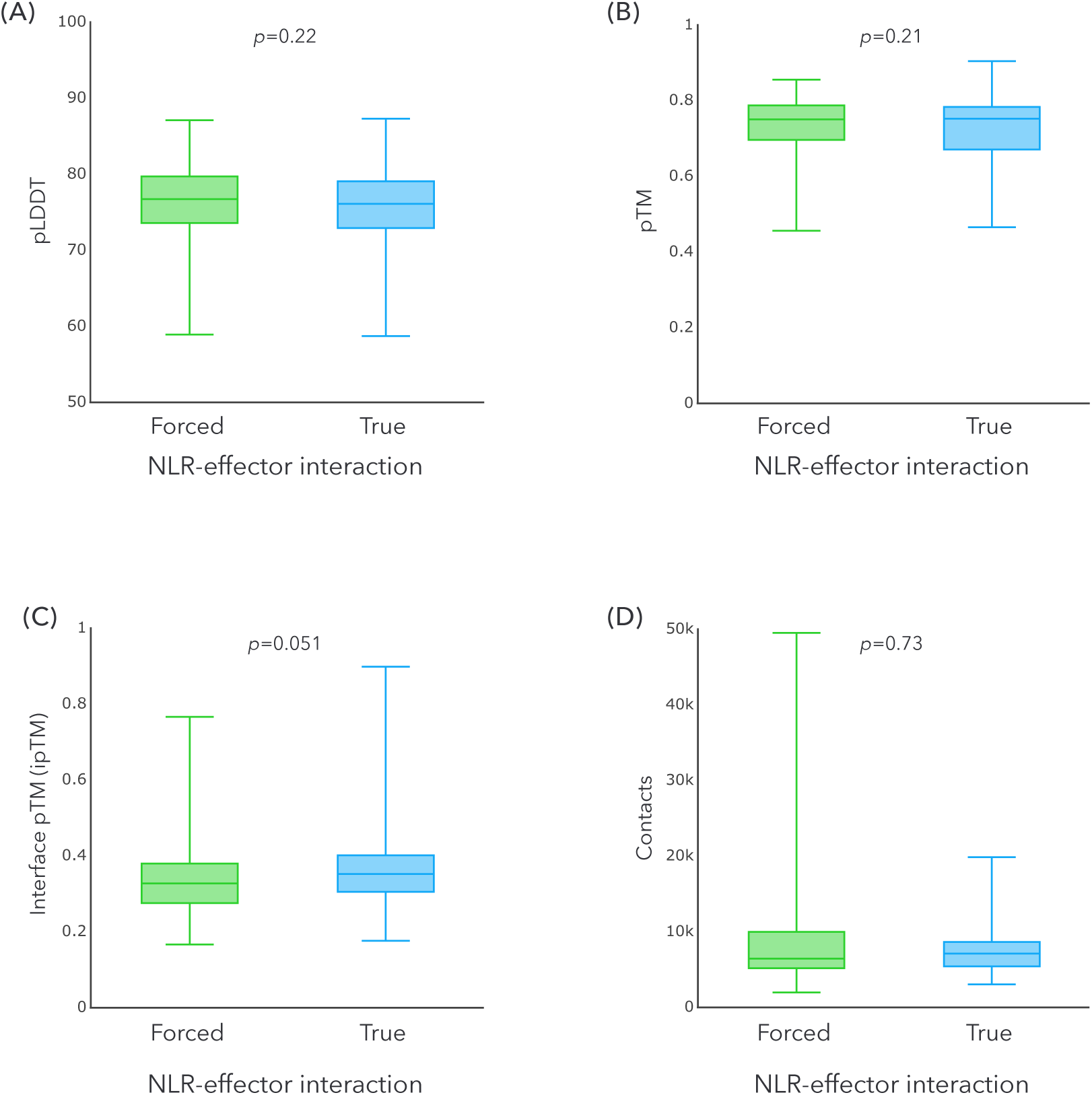
Complex and interaction interface confidence values for true and forced NLR^LRR^-effector structures predicted by AlphaFold2-Multimer. No significant differences were observed for **(A)** pLDDT, **(B)** pTM, **(C)** ipTM, or **(D)** number of contacts when comparing 93 true and 5930 forced NLR^LRR^-effector interactions. Statistical significance was calculated using Wilcoxon rank-sum tests for box-and-whisker plots, and scatterplot correlations are based on Pearson’s correlation coefficient (R = 0.45; *p* > 0.05).

The number of contacts between NLR^LRR^ and effector proteins does not differ significantly when comparing true and forced interactions (Figure 5D). Previous studies have shown that the number of contacts correlates with BA, suggesting that more contacts may be present in true NLR^LRR^-effector interactions (Raucci et al., 2018; Vangone and Bonvin, 2015). However, the specific AAs that form these contacts also influence BA (Yi et al., 2024). Thus, even though the number of predicted contacts does not differ between true and forced NLR^LRR^-effector interactions, the specific AAs involved in the interaction may contribute to differences in BA. Future research is needed to address this aspect.

### BA– and BE values for true and forced NLR^LRR^-effector complexes

Removal of predicted structures with AF confidence scores < 0.42 resulted in 58 true and 2427 forced NLR^LRR^-effector complex structures being retained for further analysis. Using the Area-Affinity web tool, the BE and BA were predicted for each NLR^LRR^-effector complex (Yang et al., 2023). As expected, there was significant variation in both BE and BA values among the NLR^LRR^-effector structures when evaluated using the combined 97 models employed by Area-Affinity (Figure 6 and Figure 7, respectively). BE values ranged between –0.049 kcal/mol (protein-protein Generated nonlinear model (linear fitting) 8) to –40.283 kcal/mol (protein-protein Linear model 10), while BA values ranged from –0.036 log(K) to –29.535 log(K) (same models, respectively). No significant difference in BE and BA values was observed between true and forced NLR^LRR^-effector interactions for a specific model. Interestingly, most models consistently predicted higher BE and BA values for some forced NLR^LRR^-effector interactions. For example, the average predicted BE and BA values across all models for the true interaction of Rpi-blb2^LRR^-Avrblb2 was –12.681 kcal/mol and –9.344 log(K), respectively, while values for the forced interaction Rpi-blb1^LRR^-Avrblb2 were – 14.728 kcal/mol and –10.798 log(K), respectively.

**Figure 6.**
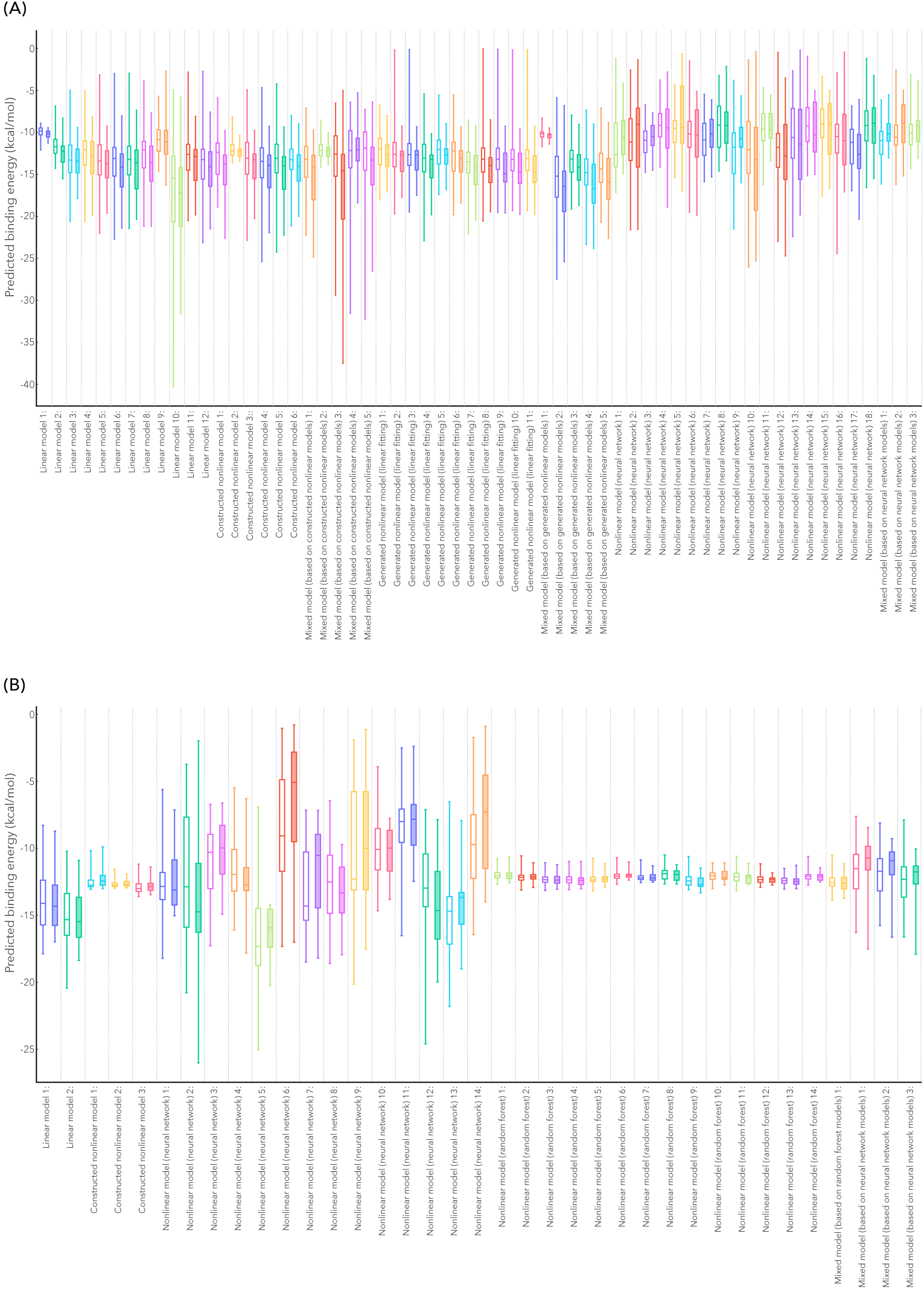
Predicted binding energies for all true and forced NLR^LRR^-effector interactions. (**A**) Binding energies were predicted for protein complexes using 60 protein-protein machine learning models, and **(B)** 37 antibody-protein machine learning models from Area-Affinity. AlphaFold2-Multimer was used for NLRLRR-effector complex prediction, with only models having AlphaFold confidence scores (0.2*pTM+0.8*ipTM) > 0.42 included in the binding energy analysis. True interactions are represented by filled box-and-whisker plots, while forced interactions are represented by unfilled plots.

**Figure 7.**
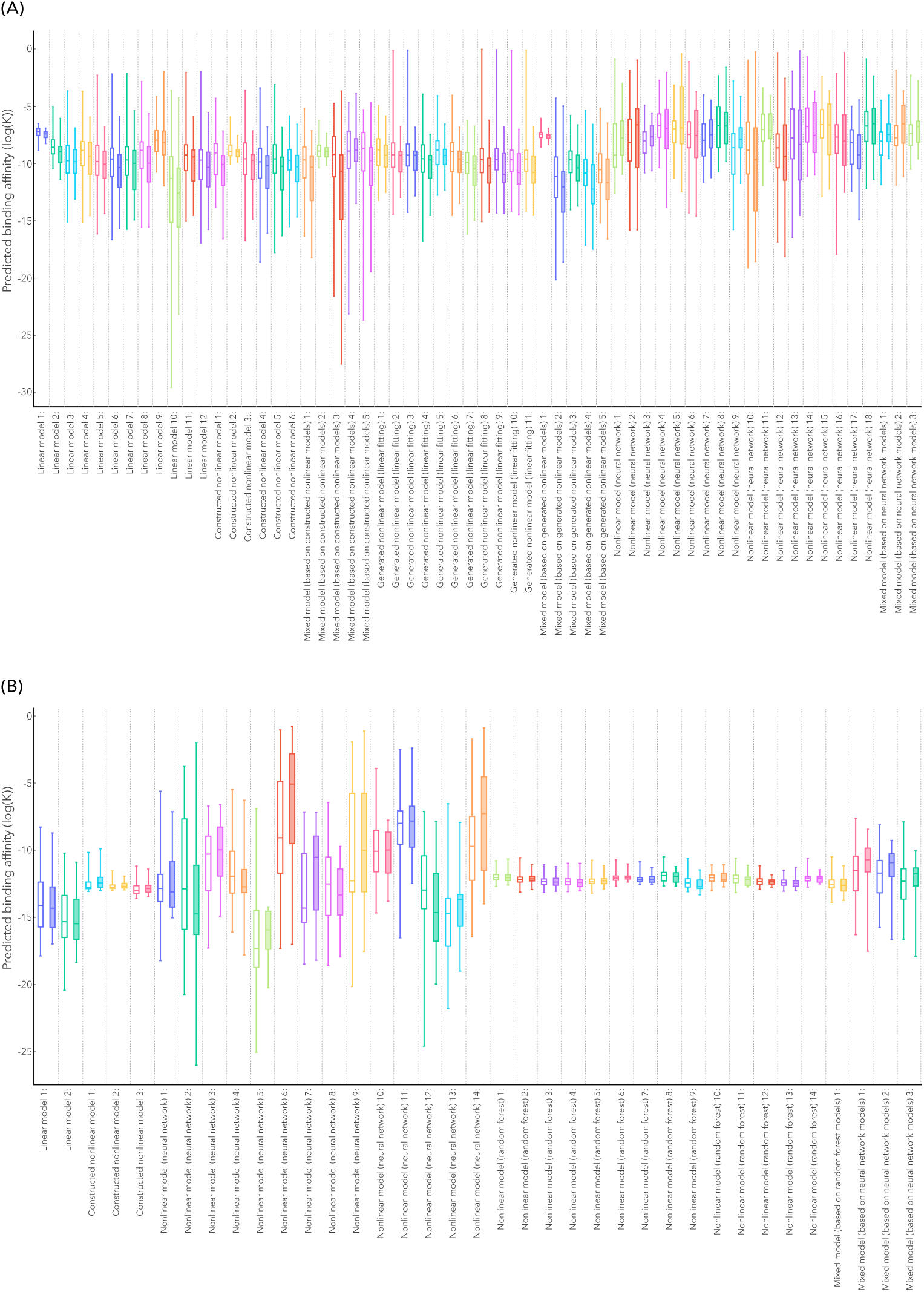
Predicted binding affinities for all true and false NLR^LRR^-effector interactions. (**A**) Binding affinities were predicted for protein complexes using 60 protein-protein machine learning models, and **(B)** 37 antibody-protein machine learning models from Area-Affinity. AlphaFold2-Multimer was used for NLR^LRR^-effector complex prediction, with only models having AlphaFold confidence scores (0.2*pTM+0.8*ipTM) > 0.42 included in the binding energy analysis. True interactions are represented by filled box-and-whisker plots, while false interactions are represented by unfilled plots.

These results suggest that the Avrblb2 effector binds more tightly to Rpi-blb1^LRR^, but this interaction does not lead to host immune response activation (Champournet et al., 2009; Oh et al.,2014). This implies that only NLR^LRR^-effector interactions with a certain BE and BA would result in NLR activation, and the subsequent immune response (McBride et al., 2022). Given that effector genes from different pathogenic species exhibit varying mutation rates, sizes, and amino acid usage, separating data based on effector type (fungal, oomycete, or bacterial), may enhance the accuracy of the Ensemble model (Erijman et al., 2014; Gómes-Pérez and Kemen, 2021; Kemen et al., 2011; Yi et al., 2024). As more NLR^LRR^-effector interaction data become available to train new Ensemble models, future studies could develop interaction-prediction programs dedicated to specific effector types, thereby improving the precision and utility of these models.

These results raise an interesting question: why do BE– and BA values show less variability for true NLR^LRR^-effector interactions when compared to forced interactions? BAs and BEs are influenced by many factors, such as the number of hydrogen bonds between interacting proteins and interface size (Kastritis and Bonvin, 2013; Klebe and Böhm, 1997). Given the diversity among true NLR^LRR^-effector interactions, larger variations in BE and BA values were expected. We hypothesize that this might be due to balancing selection, as suggested by the Red Queen model (Van Valen, 1973). According to this model, NLR^LRR^ domains are constantly evolving to ensure effector recognition, while effectors are continually evolving to avoid recognition (Rabajante et al., 2016; Råberg, 2023; Singh et al., 2018). This ongoing arms race may result in an equilibrium BA between NLRs and effectors.

Lastly, these results suggest that while effector recognition specificity is controlled by the LRR domain, the NB-ARC domain may ultimately determine NLR activation (Locci and Parker, 2024). This aligns with a previous study that observed comparable Shannon entropy scores for regions within the NB-ARC– and LRR domains, indicating equal mutation rates (Prigozhin and Krasileva, 2021). Thus, NB-ARC domains may contribute to effector binding, explaining why effectors with stronger BA to the “incorrect” LRR domain do not activate these NLRs. This also supports a hypothesis by Tamborski and Krasileva (2020) regarding NLR activation mechanisms; the authors proposed that both the “switch model” and the “equilibrium-based switch model” co-exist within a system. The “switch model” postulates that only after an effector binds to an ADP-bound NLR does an intramolecular signal induce a conformational change, allowing for an ADP-ATP switch, which then triggers NLR and defense signaling (Maekawa et al., 2011; Tameling et al., 2002; Williams et al., 2011). In contrast, the “equilibrium-based switch model” suggests that NLRs cycle between ADP– and ATP-bound states, with effectors capable of binding to either state. Together, these models imply that the NB-ARC domain is central to NLR regulation and activation. Its function may be influenced by the overall auto-inhibitory strength of other domains (CC/TIR or NB domain), or by NLR^LRR^-effector recognition.

### Implementation of the Ensemble learning model

Following the training and testing of the Ensemble learning model, a cross-validation procedure indicated a model accuracy of 99.61%. This allows NLR^LRR^-effector interactions to be classified as either being true or forced with very high accuracy. This model has been implemented as an NLR^LRR^-Effector Interaction Classification (NEIC) application, available for public use, enabling classifications for unknown interactions following the steps described in the Supplementary material. However, caution should be taken when investigating NLR^LRR^-bacterial effector interactions, as only four of the predicted NLR^LRR^-bacterial effector complexes were shown to be accurate (AF confidence score > 0.42). Consequently, the model was trained using only these four NLR^LRR^-bacterial effector complexes.

NEIC was tested on Rpi-chc1.2– and rpi-tub1.3 complexes interacting with two variations of PexRD31, following manual mutation of NLR^LRR^ sequences (Monino-Lopez et al., 2021). Results obtained from NEIC mostly correlated with experimental results from Monino-Lopez et al., (2021), as tested using co-infiltration in *N. benthamiana*, with 75% accuracy. Four mutated RB::C2 NLRs (RB::C2_14-19; RB::C2_19-A; RB::C2_19-B; RB::C2_18) were experimentally shown to activate localized cell death following PexRD31-B and PexRD31-C recognition, while four other mutants (RB::C2_19-C; RB::C2_17+18; RB::C2_16; RB::C2_16+19) failed to activate cell death responses when co-expressed with PexRD31-B and PexRD31-C. NEIC correctly classified 12/16 interactions as being true or forced. These results indicate that NEIC can be employed to investigate the effect of single AA mutations on NLR^LRR^-effector interactions, however with lower accuracy. Yet, as most interactions were indeed correctly classified, the results from NEIC may still provide a more targeted approach for the experimental validation of NLR-effector interactions. Furthermore, future studies could produce all possible NLR^LRR^ variations through *in silico* mutation, enabling the identification of crop varieties with increased resistance levels.

## Conclusion

Despite the limitations of predictions, including the reliance on AlphaFold-predicted structures, plausible hypotheses can be inferred and applied to study the underlying mechanisms of NLR-effector interactions. These hypotheses can help design and streamline experimental procedures, accelerating molecular efforts to unravel plant immune responses. We show that NLR^LRR^-effector structures with AlphaFold confidence scores > 0.42 have acceptable comparability with experimentally validated structures and can be used to investigate NLR^LRR^-effector interactions. Furthermore, we identified that BEs and BAs for true NLR^LRR^-effector interactions fall within a certain range, which can aid in identifying new interactions. All programs used in this study are freely available online, allowing the prediction of new NLR-effector interactions without the need for powerful computers or servers This provides a more targeted approach when studying these interactions *in planta*. As more data on these interactions become available, dedicated AI programs can be developed to predict these interactions with greater accuracy compared to currently available programs.

## Acknowledgments

The authors would like to express their gratitude to Prof. Warren du Plessis for his invaluable discussions regarding classification models, and the implementation thereof. Furthermore, we would like to thank Mr. Daniel Opperman for his insight on the methods used for NLR^LRR^-effector complex predictions, and comments on the manuscript.

