## Supplementary material. for "What NLR you recognizing? Predicted binding affinities- and energies may be used to identify novel NLR-effector interactions"

1. Prepare sequences for protein complex prediction
- Identify NLR- and effector domains using InterPro. This can be done using both the online web tool: https://www.ebi.ac.uk/interpro/search/sequence/
OR using a locally installed version.
- Remove signal domains from effector sequences, and CC/TIR and NB domains from NLR sequences. Thus, only use NLR LRR domain-encoding sequences.

2. Predict NLR-effector structures using AlphaFold2-Multimer
- Template mode should be set to "PDB100"
- Pair mode should be set to "Unpaired"
- Model type should be set to "alphafold_multimer-v3"

3. Identify predicted complexes with acceptable accuracy
- Calculate AlphaFold confidence scores for top ranked structures using the formula: 0.2*(pTM)+0.8*(ipTM).
- Predicted complexes with AF confidence scores ≥ 0.42 may be used for further analysis.

4. Predicting Binding energy- and affinity values
- Predict Binding energy- and affinity values using the Area-Affinity web tool: https://affinity.cuhk.edu.cn
- Values should be predicted using both the "Protein-protein", and "Antibody-antigen protein" options.
- Download results (output in .txt format)
- Combine results from "Protein-protein" and "Antibody-antigen protein" predictions. This should be done by copying- and pasting values from the "Antibody-antigen protein" .txt file (without header) directly below "Protein-protein" values.

5. Prepare NEIC input Excel file
- Copy combined Binding energy- and affinity values (without header), and paste into the "Data_Transformation.xlsx" file provided. This file orders data, with the output being in the correct format.
- Transformed data will be in the second row (coloured in blue), which can be pasted (using the "Paste special > Values" option) into a new Excel spreadsheet. This allows for binding energy- and affinity values from multiple NLR-effector complexes to be analysed by NEIC.
- The first row in the new Excel spreadsheet should contain an alphabetic header, which can be found in the first row of the "Data_Transformation.xlsx" file.
- Once the spreadsheet contains all data to be analysed, "Find and replace" all "#VALUE!" with "-1".
- Save spreadsheet in .xlsx format.

6. Install NEIC application
- Detailed instructions are available at: https://github.com/koosfick/NEIC1.0

7. Classify NLR-effector interactions using NEIC
- Import the .xlsx file prepared in step 5.
- Results will be displayed, with an interaction probability value given in the format "Forced vs Interacting". 
